# Analyzing Repast Symphony models in R with **RRepast** package

**DOI:** 10.1101/047985

**Authors:** Antonio Prestes García, Alfonso Rodríguez-Patón

**Affiliations:** Departamento de Inteligencia Artificial,Universidad Politécnica de Madrid, Campus de Montegancedo s/n, Boadilla del Monte, 28660 Madrid, Spain

## Abstract

In order to produce dependable results, the output of models must be carefully evaluated and compared to the experimental data. One of the main goals of analyzing a model is understanding the effect of input factors on the model output. This task is carried out using a methodology known as sensitivity analysis. The analysis of Individual-based Models is hindered by the lack of simple tools allowing a complete and throughout evaluation without much effort. This kind of models tends to have a high level of complexity and the manual execution of a large experimental setup is generally not a feasible choice. Thus, it is required that model evaluation should ideally be simple and robust without demanding a high level of knowledge from modelers. In this work we present the RRepast, an open source GNU R package for executing, calibrating and analyzing Repast Symphony models directly from the R environment.

## I. INTRODUCTION

The individual-based modeling is being established progressively as a main-stream and valuable tool for modeling complex processes in many distinct areas of knowledge, ranging from social science, economics to any flavor of computational and systems science such as biology, ecology and so on [1].The reason is, amongst other things, the relative ease with which detailed structural information can be incorporated into a model without the constraints of other methodologies [2].Nonetheless, the possibility of incorporating many details comes with the cost of models with a high complexity level, containing many rules and parameters for which the exact values are, in many cases, hard or impossible to determine experimentally, that is what is know as parameter uncertainty.

Model calibration is the task of estimate the set of values for input parameters of some simulation model which provides the best fitting to any empirical data set available for the system under study [3]. The estimation of acceptable values for the parameters of Individual-based Models and the analysis of uncertainty, requires specialized techniques which are complex and computationally demanding. One of the objectives of these methods are understand the relative impact of input parameters on the overall model outcomes. According to [4] most of Individual-based models published tends to omit the systematic calibration and sensitivity analysis tasks, chiefly due to the fact that modelers practitioners do not have the specific knowledge to implement or simply use the required methods. Therefore, it seems to be clear, that the availability of simple and user friendly tools for experiment design and analysis would help modelers to improve the formal quality of their models.

The Repast Symphony framework is a fast and flexible java-based environment with some built-in facilities for batching and parameter sweeping [5], widely used in many fields for building individual-based simulation models ([6], [7], [8]). Repast also has support for running GNU R ([9], [10]) from inside the framework user interface but until now it has not feasible to run Repast models from R environment for controlling model in order to implement experimental designs, parameter calibration and sensitivity analysis, therefore hindering a throughout and comprehensive verification of Individual-based models.

In addition, the real value of a computational model depends much on the ability of other researchers to reproduce and enhance the results elsewhere; in other words results must be reproducible. Hence, in order to achieve reproducibility, research methods should be stated clearly and should preferentially being backed by standard methods and software tools. In the following sections we will describe the **RRepast** package functionalities, the most significant API elements, as well as a worked example for illustrating the basic use case of the package.

## II. THE RRepast PACKAGE

The **RRepast**^1^ is an ongoing open source project developed primarily for invoking Repast Symphony models from inside GNU R environment. Additionally the package has other features for making the analysis of Individual-based models developed with Repast, extremely straightforward.

The package has two main groups of functions: the first, directly related to the integration of Repast Symphony with R, allowing the instantiation, execution and control of a model execution, as well as, gathering model output generated by any aggregated dataset defined into Repast model [11]. The second group of features exposes a complete set of methods for parameter calibration and for performing sensitivity analysis methods without much effort, also including functions for most common experimental design setups.

The first group of methods, are in turn subdivided into low and high level calls. The first type of them are the functions prefixed with the [Engine] keyword which wraps the calls to the java subsystem using the rJava package [12]. These functions are not intended for general use, instead the users should the high level calls which include, the calls depicted in the Table I.

**Table I:** The basic RRepast API Functions. These functions are used for loading, modifying the default parameters defined for model and for running the simulation.

| Function name | Description |
| --- | --- |
| **Model(d, t, o, l)** | This function creates an object instance for linking the Repast model to an R object. The required parameters are the directory where the model has been installed (*d*), the duration of simulation in Repast ticks (*t*), the name of any aggregated dataset of model for draining data generated by the model simulation (*o*) and a Boolean flag (*l*) which tells the function to call the Load method. The default value is FALSE. |
| **Load(m)** | This function loads the Repast scenario from model's directory. The only required parameter (*m*) is an instance of Repast Model created with previous function. |
| **Run(m, r, s)** | The purpose of this function is to execute a single round of simulation using just one parameter set. The parameters for this function are a model instance (*m*), the number of repetitions (*r*) and a collection of random seeds (*s*) to be used for each one of the repetitions. The only required parameter is the model instance, created with the *Model()* function. The default value for *r* is one. |
| **RunExperiment(m, r, d, F)** | Execute a complete experimental setup for different sets of parameters. The parameters required are a model instance (*m*), the number of replications (*r*), the experimental design (*d*) and finally a user provided calibration function (*F*). The experimental design parameter is an **R** data frame containing a complete set of model's parameter per row. The function returns a list with three data frame elements: the *paramset*, the *output* and *dataset* which holds respectively all simulated input parameters, the result of user provide calibration function and the complete dataset produced during the experiment execution. |
| **GetSimulationParameters(m)** | Returns the complete list of parameters declared by the model. The parameter (*m*) is an instance of Repast model generated with *Model()* call described previously. |
| **SetSimulationParameters(m, p)** | Modify several parameters at once. |
| **SaveSimulationData(t, e)** | Exports the results of Run or RunExperiment to a csv or excel files. The parameters *t* and *e* are respectively the format of exported data (xls or csv) and the experiment results returned by *RunExperiment()* |

The second group of functions inside the package contains *low level* functions for the design of experiments [13] by the user, as well as, *high level* methods which are the recommended entry point for the generation of experiments with the model. All of these *high level* functions have their names prefixed with the "Easy" keyword. The *Easy API* are designed to perform a complete and complex task with just one function call. Some of these functions are shown in the Table II and the current Easy API methods are presented in Table III.

**Table II:** The Experimental Setup API functions. These functions are used for experimental design, parameter calibration and sensitivity analysis.

| Function name | Description |
| --- | --- |
| **AddFactor(f, l, k, b, u)** | Creates the parameter collection for the experimental setup. The function requires the data frame (*f*) where parameter will be added, if this parameter is not provided a new data frame will be created. The second parameter (*l*) is the random function used internally, the default value is *runif* which will be the valid choice in many cases, the next parameter (*k*) is the name of factor, the value provided must match some parameter defined in the repast model. The following two parameters (*b*) , (*u*) are the lower and the upper range, respectively. The function returns the updated (*f*) data frame with the new parameter. |
| **AoE.RandomSampling(n, f)** | Also known as Monte Carlo sampling, generate an experimental design based making random samplings of parameter space. The function takes two parameters, the sample size (*n*) and the factor (*f*) data frame created using **AddFactor()**. The function returns the design matrix from the provided parameters. |
| **AoE.LatinHypercube(n, f)** | Generates an experimental design using the Latin Hypercube stratified sampling technique which is a more efficient sampling scheme, in terms of model evaluations, than the pure random sampling. The parameters (*n, f*) and return values are the same already described for the function **AoE.RandomSampling()**. |
| **AoE.FullFactorial(n, f)** | Creates a factorial design where the effects of all independent variables of model are studied simultaneously which implies many more model evaluations. The parameters (*n, f*) and return values are the same already described for the function **AoE.RandomSampling()**. |
| **BuildParameterSet(d, p)** | Constructs the data frame required for executing *RunExperiment()*. The function takes two parameters: the design matrix (*d*) created with one of previous functions and the declared parameters (*p*) defined in the Repast Model with the default values retrieve using the function **GetSimulationParameters()**. The functions returns a data frame with varying and fixed parameters for the experimental setup of choice. |

**Table III:** The easy API functions. These functions are the preferred entry point for the eventual users. These "Easy" functions lump together a complete experiment task in just one call, reducing the coding needs to the minimum.

| Function name | Description |
| --- | --- |
| **Easy.Stability(d,o,t,f,s,r,v,F)** | Evaluate the behavior of model output in order to determine the minimum required number of replication of chosen experimental setup. The function accept the following parameters: the model installation directory (*d*), the aggregated data source defined within the Repast model (*o*), the simulation time in Repast ticks (*t*) which default value is 300 ticks, the input factors to be sampled (*f*) created with the previously mentioned function **AddFactor()**, the number of parameter samples (*s*), the desired number of replications to be tried (*r*) being the default value 100, the output variables of interest which will be checked for their stability and convergence of the coefficient of variation (*v*); if this parameter is leaved empty all output variables are checked and finally the user provided calibration function (*F*) for determining the best input parameter combination. |
| **Easy.Morris(d,o,t,f,p,s,r,F)** | This function performs all required tasks for carrying out the method of Morris for screening. The parameters are practically the same as described for the previous function with exception of parameters (*p*) and (*s*) which are respectively the levels of input factors and the number of sampling points of Morris method [14]. |
| **Easy.Sobol(d,o,t,f,n,r,F)** | Encapsulate all required steps for performing sensitivity analysis using Sobol method. The method of Sobol is a global sensitivity analysis technique based on the decomposition of output variance ([15], [14]). The parameter semantics are the same already described: the model installation directory (*d*), the aggregated data source defined within the Repast model (*o*), the simulation time in Repast ticks (*t*) , the input factors to be sampled (*f*), the sample size (*n*), the desired number of replications (*r*) and calibration function (*F*). |
| **Easy.Calibration(d,o,t,f,n,r,F)** | This function estimates the best set of input parameters or factors performing a set of experiments in order to sample the calibration function. The objective of this function is to minimize the output of calibration function provided by the user. |

## III. **RRepast** IN ACTION

In this section we will provide some small examples on how to use the **RRepast** package for running Repast models and analyzing the data produced. In order to get the model running from R code, some minimal steps must be carried out before calling Repast code.

1) Build an installer and install the Repast model.
2) Add the **rrepast-integration.jar** file, included in **RRepast** distribution, to the **lib** directory of the installed Repast model.
3) Add the integration configuration to scenario file in the. rs directory of the installed model. The integration consists in the following code: *<model.initializer dass="org.haldane.rrepast.ModelImtializerBmker"/>*

Once the previous steps are completed we are ready for running the model. The minimal code to execute the model is presented in Figure 1

**Figure 1:**
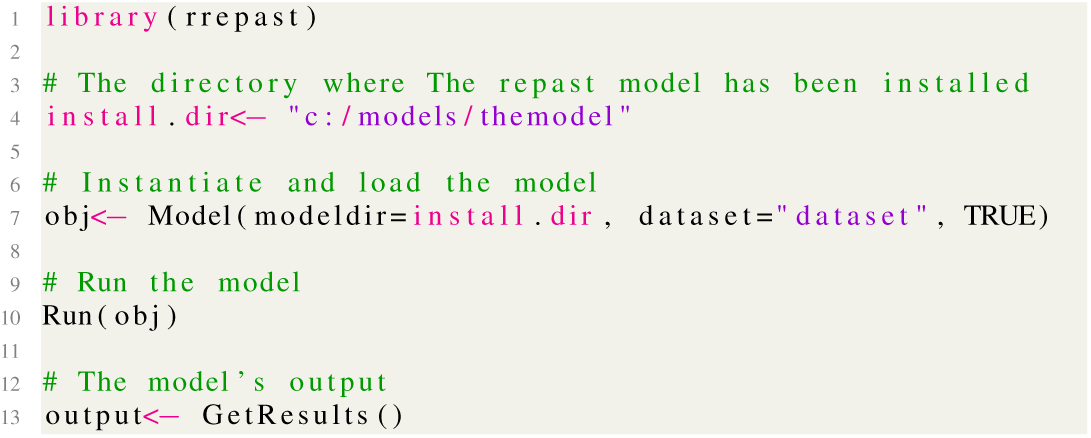
**The minimal code for running a Repast model from R. The Boolean value in** *Model()* **tells RRepast to auto load the model’s scenario.**

In addition to the basic functionality for loading and running a model and retrieving the complete output of any dataset defined in the Repast model, the package contains an implementation of common techniques for screening and global sensitivity analysis as well as for verifying the stability of output variables. These functionalities are readily accessible, requiring very few lines of code. In the simplest case the modeler only has to complete three tasks for getting the experiment done. The first one is to define a calibration function. The calibration function must return zero for the best fit and a other number greater than zero otherwise. How the criteria are implemented is up to the modeler. That function is called internally by RRepast and has a specific format. The parameters passed to the function are the current set parameter used and the complete content of Model dataset output. The function must return a **cbind()** containing all individual criteria and optionally the sum of individual criteria.

In order to provide some more realist examples we have used the BactoSIM Repast model, which is an spatially explicit individual-based model for simulating the plasmid spread on a surface attached bacterial colony [16].

The **BactoSIM** simulation model has several parameters but we want to focus just on four of them keeping all other fixed. Thus, let’s say, we want to evaluate the parameters named *gamma(), cyclePoint, conjugationCons* and *pilusExpression-Cost*. For accomplishing this task we will use the *Easy* API functions described in Table III. These functions return a list holding three elements:

- **experiment**. The experiment is also a list holding the parameter set (paramset), the calibration function output (output) and the experiment raw dataset (dataset). These three entities are connected by a column named **pset**.
- **object**. The reference to the object used which could be Morris or a Sobol instance.
- **charts**. Contains the reference to the plots generated.

Therefore, the first step could be to determine the required number of replications for the simulation experiments using the *Easy.Stability()* which output can be seen in Figure 2. The output shows on the abscissa the number of repetitions and on the ordinates the coefficient of variation for the desired output variable.

**Figure 2:**
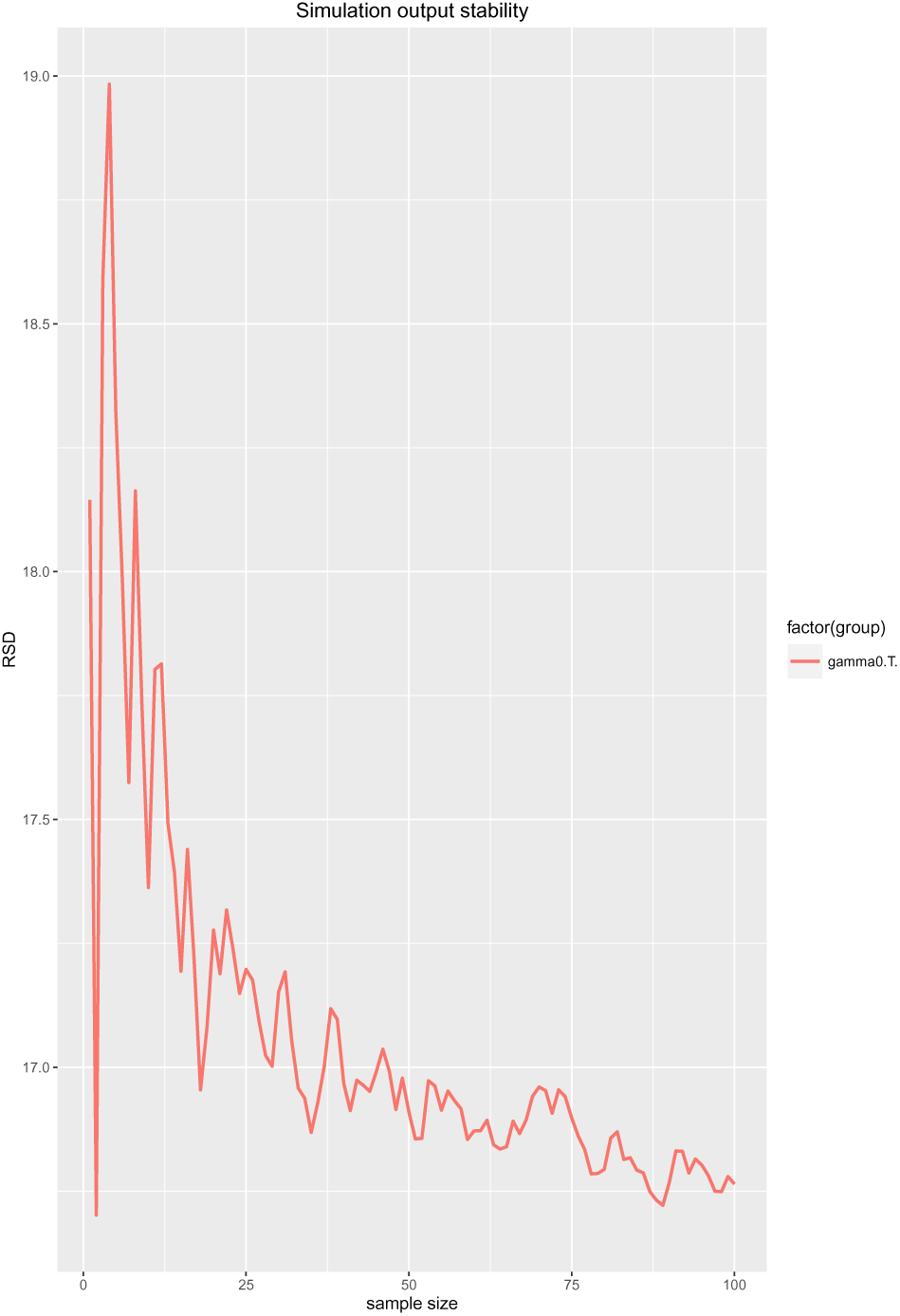
The stability of model output.It is possible to observe how, as far the number of replications of the experimental setup increases, the value of the coefficient of variation converges to a common value.

The listing shown in Figure 3 is an example of how easy is to analyze simulation experiments using RRepast. That is all code required to perform the Morris screening method for the BactoSIM Model. One of the outputs of Morris method is presented in Figure 4.

**Figure 3:**
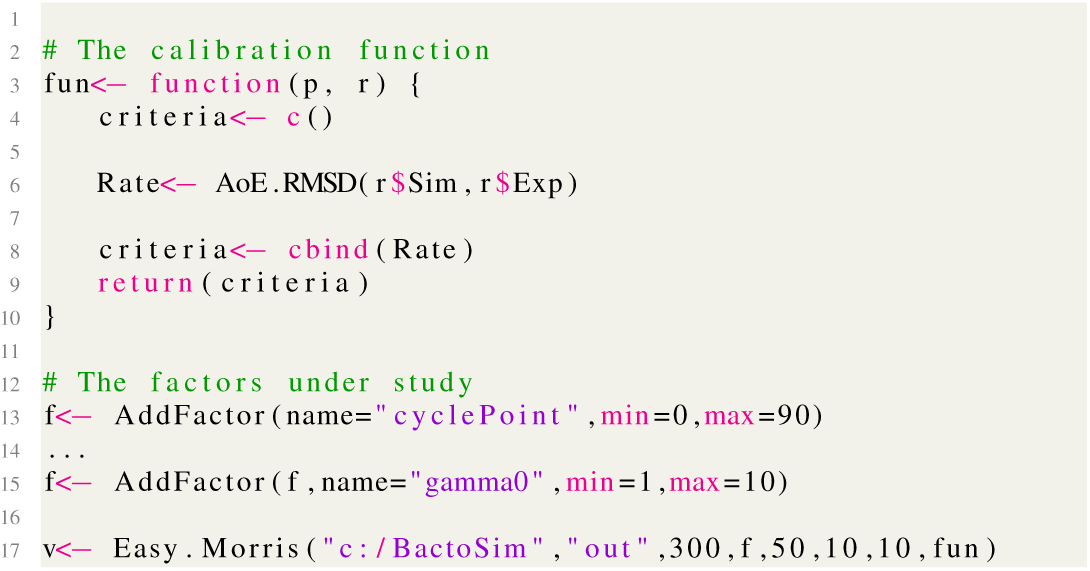
The complete listing for perform the Morris’s screening method. In the line 6 we define the **Rate** calibration criteria which is root-mean-square deviation between simulated and observed values. In lines 13 to 15 we create the input factor collection with their range of variation and finally line 17 shows the call of Easy.Morris function.

**Figure 4:**
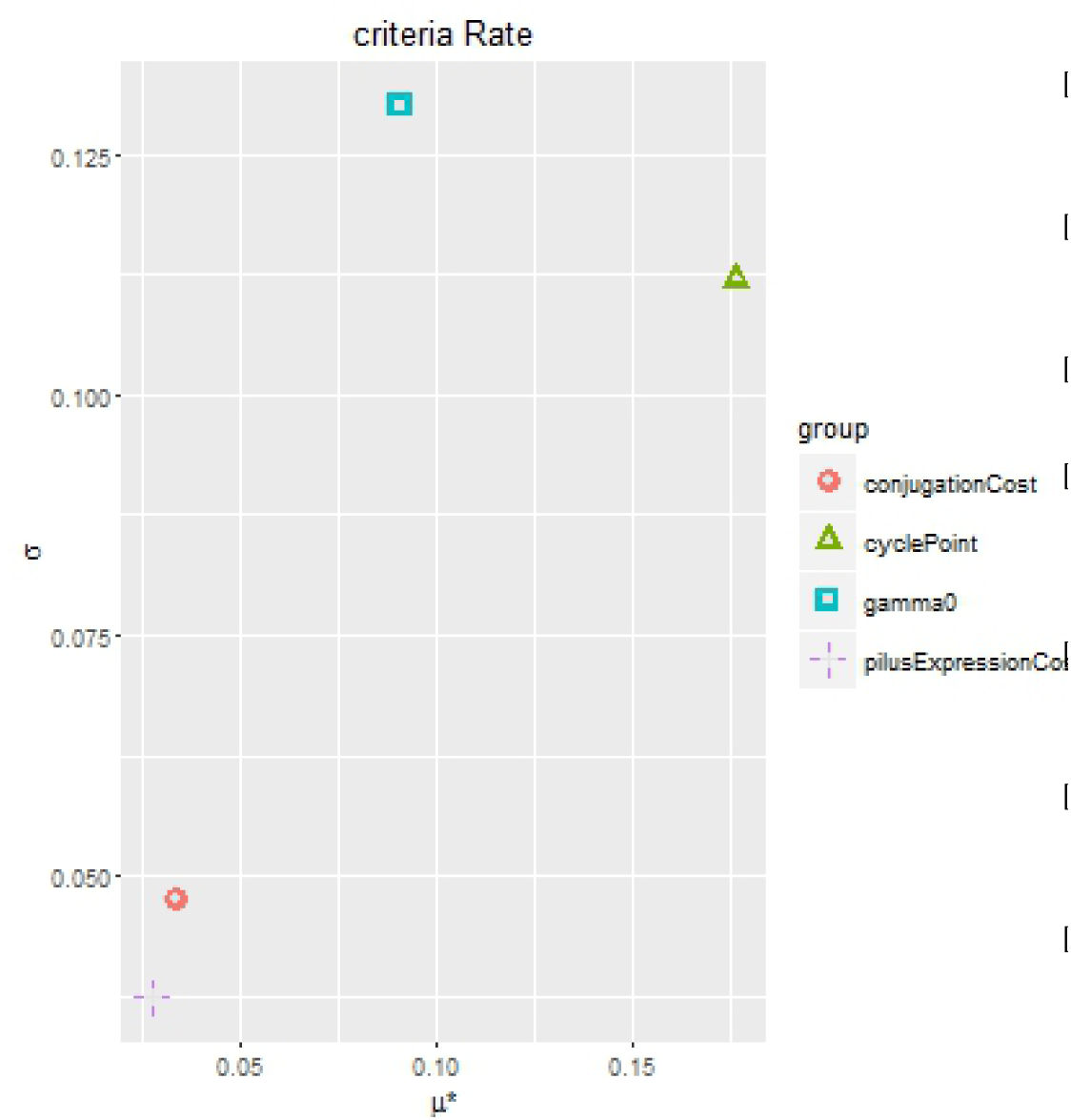
One of the output charts for Morris’s screening method.The chart shows that the most import parameter for the Rate calibration metric is the cyclePoint followed by the gammaO.

Finally we could decide, using the output of Morris method, to discard some of the parameters and focus only on those more important to perform the Sobol method. One of the output charts of Sobol method showing the indices and the confidence interval are show in Figure 5.

**Figure 5:**
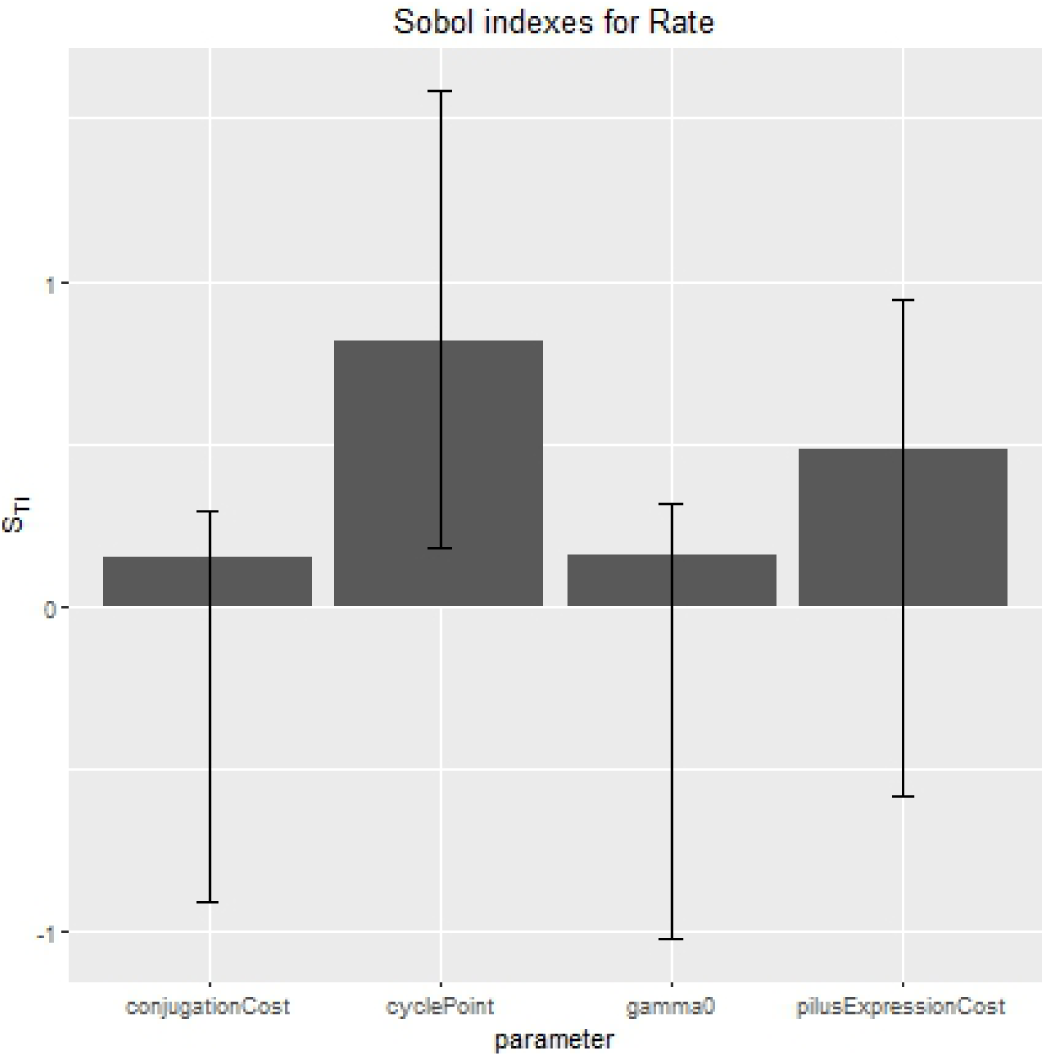
The output chart for Sobol method.The Sobol output shows that the dominant parameter is the cyclePoint but differently from Morris method the second in importance seems to be the pilusExpressionCost.

## IV. CONCLUSIONS

In this report we have presented the basic aspects of **RRepast** package and how it could be used for perform the basic experimental setup of Repast Models. The API functions shown here are planned to be stable but they are not frozen yet as the project is still a work in progress, hence some slight variations may occur from version to version. Future versions will include more out-of-box functions for the statistical analysis of the model output and we are also evaluating the possibility of parallelize the multiple model’s executions required by the sensitivity analysis methodologies.

One of main drawback of analyzing individual-based models is the computational cost and the time required to complete an experimental setup for any model with a medium complexity level and a high number of agents being simulated. The simulations are safe and relatively easy to distribute as the same code will be executed for a different set of parameters but there are no need to communicate instances of experimental setup. Recently some interest has been shown on using Docker container technology for scientific research [17] and we are exploring that technology for easy deployment of the model execution across many nodes seamlessly.

## ACKNOWLEDGMENTS

This work was supported by the European FP7 - ICT - FET EU research project: 612146 (PLASWIRES "Plasmids as Wires" project) www.plaswires.eu and by Spanish Government (MINECO) research grant TIN2012-36992.

## Footnotes

1 The software can be found on the following CRAN URL: https://cran.r-project.org/web/packages/rrepast/

